# Statistical Integration of Two Omics Datasets Using GO2PLS

**DOI:** 10.1101/2020.08.31.274175

**Authors:** Zhujie Gu, Said el Bouhaddani, Jiayi Pei, Jeanine Houwing-Duistermaat, Hae-Won Uh

## Abstract

**Background:** Nowadays, multiple omics data are measured on the same samples in the belief that these different omics datasets represent various aspects of the underlying biological systems. Integrating these omics datasets will facilitate the understanding of the systems. For this purpose, various methods have been proposed, such as Partial Least Squares (PLS), decomposing two datasets into joint and residual subspaces. Since omics data are heterogeneous, the joint components in PLS will contain variation specific to each dataset. To account for this, Two-way Orthogonal Partial Least Squares (O2PLS) captures the heterogeneity by introducing orthogonal subspaces and better estimates the joint subspaces. However, the latent components spanning the joint subspaces in O2PLS are linear combinations of all variables, while it might be of interest to identify a small subset relevant to the research question. To obtain sparsity, we extend O2PLS to Group Sparse O2PLS (GO2PLS) that utilizes biological information on group structures among variables and performs group selection in the joint subspace.

**Results:** The simulation study showed that introducing sparsity improved the feature selection performance. Furthermore, incorporating group structures increased robustness of the feature selection procedure. GO2PLS performed optimally in terms of accuracy of joint score estimation, joint loading estimation, and feature selection. We applied GO2PLS to datasets from two studies: TwinsUK (a population study) and CVON-DOSIS (a small case-control study). In the first, we incorporated biological information on the group structures of the methylation CpG sites when integrating the methylation dataset with the IgG glycomics data. The targeted genes of the selected methylation groups turned out to be relevant to the immune system, in which the IgG glycans play important roles. In the second, we selected regulatory regions and transcripts that explained the covariance between regulomics and transcriptomics data. The corresponding genes of the selected features appeared to be relevant to heart muscle disease.

**Conclusions:** GO2PLS integrates two omics datasets to help understand the underlying system that involves both omics levels. It incorporates external >group information and performs group selection, resulting in a small subset of features that best explain the relationship between two omics datasets for better interpretability.

## Background

With the advancements in high throughput technology, multiple omics data are commonly available on the same subjects. To identify a set of relevant related features across the omics levels, these datasets need to be integrated and analyzed jointly. For statistical integration of omics data, there are several challenges to overcome: complex correlation structure within and between omics data, high-dimensionality (*p* ≫ *n*, or “large *p*, small *n*”), heterogeneity between different omics datasets, and selection of relevant features in each dataset. To deal with the first two challenges, Partial Least Squares (PLS) has been proposed [5, 30]. Dimension reduction is achieved by decomposing two datasets *X* and *Y* into joint and residual subspaces. The joint (low-dimensional) subspace of one dataset represents the best approximation of *X* or *Y* based on maximizing the covariance of the two. However, by integrating two heterogeneous omics datasets, the PLS joint components also contain (strong) omic-specific variation. This heterogeneity can be caused by differences (e.g. between methylation and glycomics) in size, distribution, and measurement platform. Ignoring these omic-specific characteristics (variation specific to each of the data) in the model may lead to a biased representation of the underlying system. Two-way orthogonal partial least squares (O2PLS) [24, 9] was proposed to decompose each dataset into joint, orthogonal, and residual subspaces. The orthogonal subspaces in *X* and *Y* capture variation unrelated to each other, making the joint subspaces better estimates for the true relation between X and Y. Hence, O2PLS accounts for the heterogeneity of two omics datasets. However, the resulting low-dimensional latent components spanning the joint subspaces are linear combinations of all the observed variables. Therefore, to select a small subset of relevant features for better interpretation, one can impose sparsity on the loadings of the principal components. A straightforward approach is to ignore all loadings smaller than some threshold value, effectively treating them as zero, which can be misleading [13].

Several sparse methods based on PLS have been proposed. Chun and Keleş proposed sparse PLS (SPLS) [8] which fits PLS on a reduced *X* space, consisting of pre-selected *X*-variables using a penalized regression. Sparse PLS (sPLS) by Lê Cao et al. [15] imposes *L*_1_ penalty on the singular value decomposition (SVD) of the covariance matrix of *X* and *Y*, resulting in sparse loading vectors for both datasets. Often it is of interest to select a group of features instead of individual features, e.g. features within a gene or a pathway. By so doing, one can improve power by identifying aggregate effects of the selected features [25, 31, 16]. Liquet et al. extended sPLS to group PLS (gPLS) [16], imposing group-wise *L*_2_ penalties on the loadings of the pre-defined feature groups. It results in group-wise sparsity (i.e., features belonging to the same group will always be selected altogether).

In this work, we propose to extend O2PLS to incorporate sparsity, called Group Sparse O2PLS (GO2PLS). GO2PLS obtains sparse solutions by pushing a large number of small non-zero weights (or loading values) to zeros, instead of employing hard thresholding using arbitrary cut-off values. Therefore, GO2PLS constructs joint low-dimensional latent components representing the underlying systems involving both omics levels while taking into account the heterogeneity of different omics data, incorporates external biological information such as known group structure, and performs variable selection by imposing group-wise penalties on the loading vectors in the joint subspaces.

For illustration, we apply GO2PLS to datasets from two studies. Firstly, TwinsUK is a population based study [19, 17], where methylation (482K CpG sites) and 22 immunoglobulin G (IgG) glycans were measured. A previous research [27] suggested the presence of an indirect influence of methylation on IgG glycosylation that may in part capture environmental exposures. We integrate the two omics datasets, aiming to identify genes of CpG sites affecting IgG glycosylation. In the CVON-DOSIS case-control study [1], regulomics (histone modification) and transcriptomics data were measured on 13 hypertrophic cardiomyopathy (HCM) patients and 10 controls. Histone modification can have an impact on gene expression. Therefore we integrate the two omics datasets and identify a small set of regulatory regions and transcripts explaining this relationship. Moreover, the extreme imbalance in a high-dimensional setting (33K ChIP-seq and 15K RNA-seq vs 23 subjects) poses computational challenges. The resulting selected features are further studied using gene set enrichment analysis [21]. Several possible scenarios containing these characteristics are designed and investigated in an extensive simulation study.

This paper is organized as follows. In the methods section, an overview of O2PLS is presented, followed by the formulation of GO2PLS. Via a simulation study, we explore the properties of GO2PLS and compare its performance to other competitive methods. We then apply GO2PLS to integrate methylation and glycomics in the TwinsUK study and regulomics and transcriptomics in the CVON-DOSIS study. We conclude with a discussion and possible directions to further extend the method.

## Methods

### Data description

#### TwinsUK datasets

Whole blood methylation (using Infinium HumanMethylation450 BeadChip) and IgG glycomics (Ultra Performance Liquid Chromatography) data were measured on 405 independent individuals, among which 392 are females and 13 are males. The age ranges from 18 to 81, with a median of 58. The methylation dataset consists of beta values (ratio of intensities between methylated and unmethylated alleles) at 482563 CpG sites. CpG sites with missing values, on allosomes, or labeled cross-active [7] were removed. We kept only the CpG sites on CpG islands or surrounding areas (shelves and shores) that mapped to genetic regions. Age, sex, batch effect, and cell counts were corrected for using multiple regression. The glycomics dataset contains 22 glycan peaks. These peaks were normalized using median quotient (MQ) normalization [26], log-transformed, and adjusted for batch effect, age, and sex as well. The remaining 126299 CpG sites were then divided into 16892 groups based on their target genes (biological information from the UCSC database [14, 2]). No group information was available for the glycomics data.

#### CVON-DOSIS datasets

In the CVON-DOSIS study, regulomics and transcriptomics datasets were measured on the samples taken from the heart tissues of 13 HCM patients and 10 healthy controls. HCM is a heart muscle disease that makes it harder for the heart to pump blood, leading to heart failure. The regulomics data were measured using ChIP-seq, providing counts of histone modification H3K27ac in 33642 regulatory regions. The transcriptomics data contain counts of 15882 transcripts, measured by RNA-seq. The raw counts of regulomics data were normalized with reads per kilobase million (RPKM) to adjust for sequencing depth. Transcriptomics data were normalized with counts per million (CPM) with effective library size (estimated using the TMM method in EdgeR R package [18]). Further, both normalized data were log-transformed.

### Two-way Orthogonal Partial Least Squares (O2PLS)

let *X* and *Y* be two data matrices with the number of rows equal to the sample size *N* and the number of columns equal to the dimensionality *p* and *q*, respectively. Let the number of joint, *X*-orthogonal (unrelated to *Y*) and *Y* -orthogonal components be *K*, *K*_*x*_ and *K*_*y*_, respectively, where *K*, *K*_*x*_ and *K*_*y*_ are typically much smaller than *p* and *q*. The O2PLS model decomposes *X* and *Y* as follows:

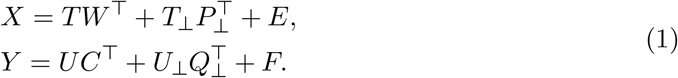

The relation between *X* and *Y* is captured through the inner relation between T and U,

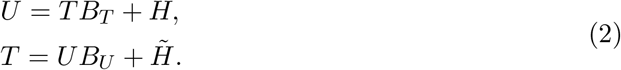

In this model, the scores are: *T* (*N* × *K*), *U* (*N* × *K*), *T*_⊥_ (*N* × *K*_*x*_), *U*_⊥_ (*N* × *K*_*y*_). They represent projections of the observed data *X* and *Y* to lower-dimensional subspaces. The loadings, *W* (*p* × *K*), *C* (*q* × *K*), *P*_⊥_ (*p* × *K*_*x*_), *Q*_⊥_ (*q* × *K*_*y*_), indicate relative importance of each X and Y variable in forming the corresponding scores. Further, *E* (*N* × *p*), *F* (*N* × *q*), *H* (*N* × *K*), 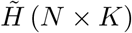, represent the residual matrices.

In O2PLS, estimates of the joint subspaces are obtained by first filtering out the orthogonal variation. The filtered data matrices 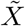 and 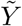 are constructed as follows:

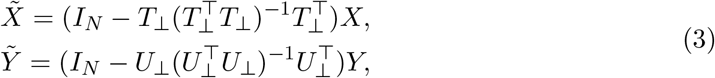

where *T*_⊥_ *U*_⊥_ are estimates for the orthogonal subspaces, and *I*_*N*_ is identity matrix of size *N*. For more details see [24]. The joint parts maximize the covariance between the joint scores 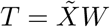 and 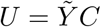. Here, *W* and *C* consist of loading vectors (*w*_1_,…, *w*_*K*_) and (*c*_1_,…, *c*_*K*_), which can be found as the right and left singular vectors of the covariance matrix 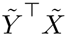 [9]. Calculating and storing 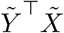 of dimension *q* × *p* can be cumbersome for high dimensional omics data. Therefore we consider the following optimization problem sequentially for components *k* = 1,…, *K*:

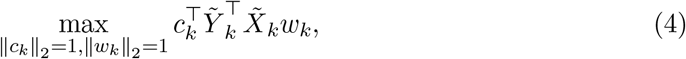

where parameters *w*_*k*_, *c*_*k*_ are the loading vectors of the *k*-th joint components and 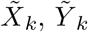 are the filtered data matrices after *k* − 1 times of deflation. This can be solved efficiently using NIPALS [29] algorithm, which starts with random initialization of the *X*-space score vector *t* and repeats a sequence of the following steps until convergence:

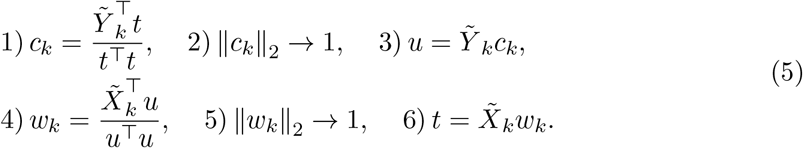

In step 1 and 4, *Y*_*k*_ and *X*_*k*_ are projected onto the *X*-space score vector *t* and the *Y* -space score *u* to get the loading vectors *c*_*k*_ and *w*_*k*_. The loading vectors are then unitized (step 2 and 5) and used to calculated the new scores *u* and *t*. Convergence of the algorithm is guaranteed. A detailed description and proof of optimality of the O2PLS algorithm can be found in [24, 9].

While standard cross-validation (CV) over a 3-dimensional grid is often used to determine the optimal number of components *K*, *K*_*x*_, and *K*_*y*_, the procedure is not optimal for O2PLS, since there is not a single optimization criterion for all three parameters. As in [9], we use an alternative CV procedure that first performs a 2-dimensional grid search of *K*_*x*_ and *K*_*y*_, with a fixed *K*, to optimize prediction performance of *T* → *U* and *U* → *T*. Then a sequential search of optimal *K* is conducted to minimize the sum of mean squared errors (MSE) of prediction concerning *X* → *Y* and *Y* → *X*.

### Group Sparse O2PLS (GO2PLS)

GO2PLS extends O2PLS by introducing a penalty in the NIPALS optimization on the filtered data 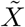 and 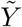. This penalty encourages sparse, or group-sparse solutions for the joint loading matrices *W* and *C*, leading to a subset of the original features corresponding to non-zero loading values being selected in each joint component.

Briefly, we introduce an *L*_1_ penalty on each pair of joint loading vectors. The optimization problem for the *k*-th pair of joint loadings *c*_*k*_, *w*_*k*_ is:

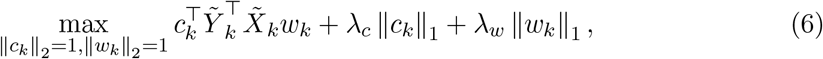

where *λ*_*c*_, *λ*_*w*_ are penalization parameters that regulate the sparsity level. The optimization problem (6) can be solved [28] by iterating over the *k*-th pair of joint loadings,

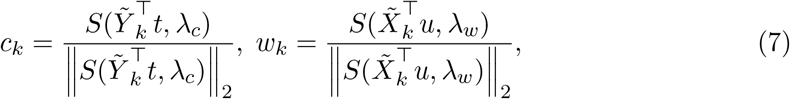

where 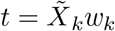 and 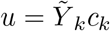. Here, *S*(·) is the soft thresholding operator: *S*(*a,* const) = *sgn*(*a*)(|*a*| − const)_+_ (const ≥ 0 is a non-negative constant, (*x*)_+_ equals to *x* if *x* > 0 and equals to 0 if *x* ≤ 0).

To perform group selection, we impose group-wise *L*_2_ penalty on the joint loading vectors. Let 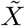 and 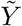 be partitioned into *J* (*J* ≤ *p*) and *M* (*M* ≤ *q*) groups, respectively. The submatrices 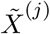 and 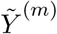 (*j* = 1,…, *J*; *m* = 1,…, *M*) contain the *j*-th and *m*-th group of variables, with corresponding loading vectors *w*^(*j*)^ (of size *p*_*j*_) and *c*^(*m*)^ (of size *q*_*m*_). The optimization problem for the *k*-th pair of loading vectors 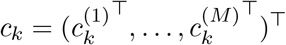 and 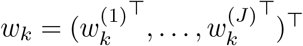 can be written as follows:

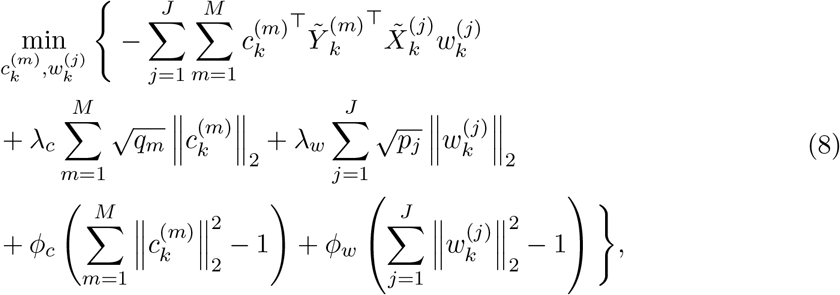

where the last two terms are reformulations of the unit norm constraints on *c*_*k*_ and *w*_*k*_, with *ϕ*_*c*_ and *ϕ*_*w*_ being the Lagrangian multipliers. The effective penalization parameters on each group (*λ*_*c*_, *λ*_*w*_) are adjusted by the square root of the group size to correct for the fact that larger groups are more likely to be selected. This optimization problem can be solved using block coordinate descent (for details, see Additional file 1). The solution takes the form:

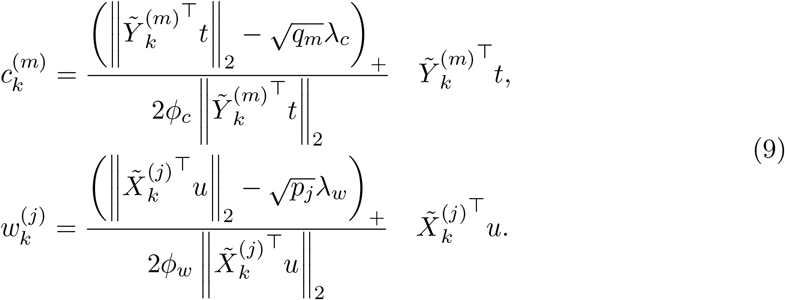

The 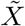-variables within the *j*-th group will have non-zero weights if 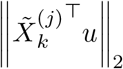 (i.e., the contribution of the whole group to the covariance) is larger than the size-adjusted penalization parameter 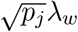. In the same way, the 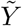-variables within the *m*-th group will be assigned non-zero loading values if 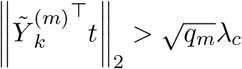.

Note that when all the groups have size 1, the summation of group-wise *L*_2_ penalties is equivalent to an *L*_1_ penalty on the unpartitioned loading vector and individual features will be selected (i.e., (8) reduces to (6)). In this specific case, to avoid confusion, we call the method Sparse O2PLS (SO2PLS). When the penalization parameters *λ*_*w*_ = *λ*_*c*_ = 0, GO2PLS becomes to O2PLS. If the number of orthogonal components *K*_*x*_ = *K*_*y*_ = 0, GO2PLS, SO2PLS, O2PLS are equivalent to gPLS, sPLS, and PLS, respectively.

The *k*-th pair of joint loadings are orthogonalized with respect to the previous *k* − 1 loading vectors. Let *π* be an index set for selected variables in *w*_*k*_. The orthogonalization is achieved by first projecting 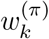 onto 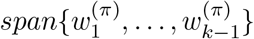, and then subtracting this projection from 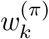. When the previous *k* − 1 components do not select any variable in *π*, 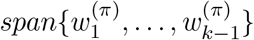 is actually a zero subspace and no orthogonalization is needed.

To determine the optimal sparsity level, it is more convenient and intuitive to focus on the number of selected 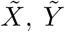 groups (donote *h*_*x*_, *h*_*y*_, respectively). If prior biological knowledge does not already specify certain *h*_*x*_ and *h*_*y*_, cross-validation can be used to search for combinations of *h*_*x*_ and *h*_*y*_ that maximize the covariance between each pair of estimated joint components 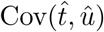. Similar to LASSO [22], the “one-standard-error-rule” [12] can be applied to obtain a more stable CV result. The GO2PLS algorithm is described below:

**Algorithm.**
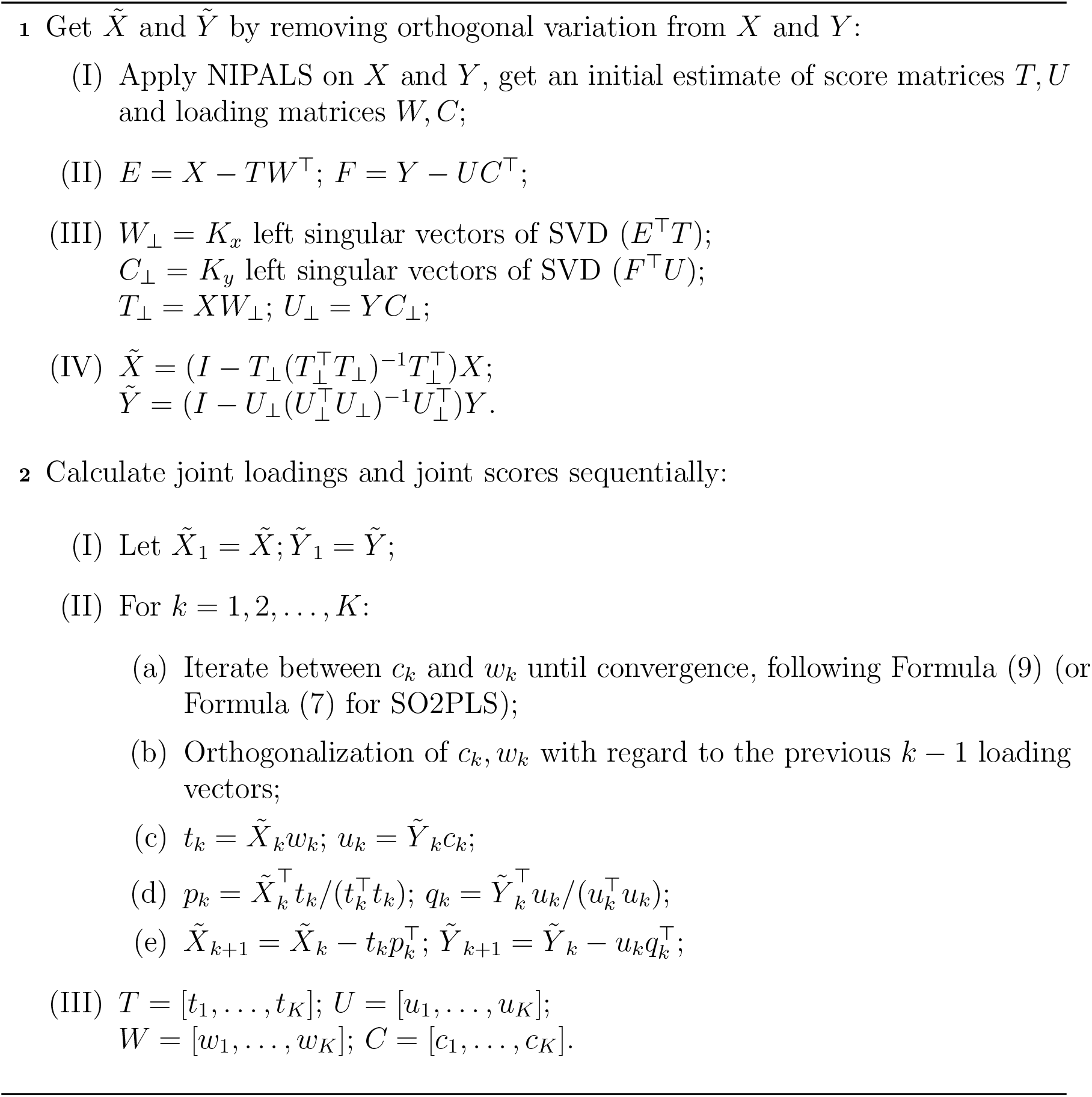
GO2PLS

### Simulation Study

We evaluate the performance of GO2PLS in two scenarios. First, we investigate the ability to select the relevant groups under various scenarios, focusing on the joint subspace, where the group selection takes place. Second, we compare the performance of GO2PLS and SO2PLS with other methods: O2PLS, PLS, sPLS, and gPLS. We investigate joint score estimation, joint loading estimation, and feature selection performances.

In the first scenario, we set the number of variables in *X* and *Y* to be *p* = 5000 and *q* = 20, respectively. There are 10 groups of variables in *X* with non-zero loading values. The first 5 groups have group sizes of 100, 50, 20, 5, and 1, respectively, in which all the variables have loading values equal to 1. The remaining 5 groups are of size 10, with loading values of variables equal to 5. Note that large loading values are assigned to the latter 5 groups to make the detection of the first 5 groups more difficult. The remaining variables have zero loading values and are divided into groups of size 10. All the *Y* -variables have the same loading values and are not grouped. The sample size *N* is set to 30. We simulate both data matrices with 1 joint component (*T* and *U* from Equation 1 are both standard normally distributed and have correlation 1). We perform 1000 simulation runs and record the number of the runs GO2PLS selected relevant groups; we compute the proportion of each truly relevant group (with non-zero loadings) being selected across the simulation runs (number of times being selected divided by 1000). The group importance measurement 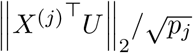, that determines whether a group is selected or not is recorded for the first 5 groups (with loading value 1) to investigate the stability of the selection procedure.

In the second scenario, we vary the sample size *N* from 30 to 600, and set *p* = 20000 and *q* = 10000, mimicking the dimensionality of the CVON-DOSIS datasets. Both *X*- and *Y* - variables are evenly divided into 1000 groups. For each joint component, we select 50 relevant groups and assign non-zero loadings to the variables contained in them. Within each group, variables have the same loading values: 1 for the first group, 2 for the second,…, and 50 for the last relevant group. We set the number of joint components *K* = 2 and the number of orthogonal components *K*_*x*_ = *K*_*y*_ = 1. The scores *T, T*_⊥_, *U, U*_⊥_ from Equation 1 are generated from normal distributions with zero mean. The relationship between the joint scores is represented by *U* = *T* + *H*, where *H* accounts for 20% of the variation in *U*. The noise matrices E, F are generated from normal distributions with zero mean and variance such that the variance of the noise matrix accounts for a proportion *α* (0 < *α* < 1) of the variance of the data matrix (i.e., *α* = Var(*E*)/Var(*X*) = Var(*F*)/Var(*Y*)). The ratio of the variance of the orthogonal components to the variance of the joint components 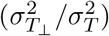, and noise level *α* are varied. For evaluating the accuracy of the joint score estimation, we computed 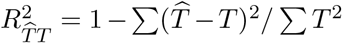 and 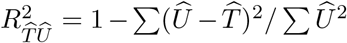, which quantify how well the true parameter *T* and the estimated *Y* -joint component 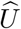 can be explained by the estimated *X*-joint component 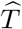. The performance of feature selection and the accuracy of estimated loadings are evaluated by true positive rate (TPR = TP/(TP+FN), where TP = True Positive, FN = False Negative) and 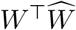, which represents the cosine of the angle between the estimated loading vector and the true one. The performances of all methods are evaluated on an independent test dataset of size 1000. For each setting, 500 replications are generated.

An overview of scenario settings is presented in Table 1, 2. To make a clearer comparison of the behavior across all the methods, we use the optimum values for the tuning parameters (number of components and number of relevant variables or groups).

**Table 1:** Settings of Scenario 1 to study the performance of selecting relevant groups.

| Measure | Selection proportion; $\frac{\ X^{(j)\top}U\ _2}{\sqrt{p_j}}$ |
| --- | --- |
| $p; q$ | 5000; 20 |
| relevant group sizes | 100; 50; 20; 5; 1 |
| $N$ | 30 |
| noise level $\alpha$ | [0.1, 0.2, 0.3, 0.4, 0.5, 0.6, 0.7, 0.8, 0.9] |

**Table 2:** Settings of Scenario 2 to compare the performances regarding joint score estimation, joint loading estimation, and feature select.

| Methods | GO2PLS; SO2PLS; O2PLS; gPLS; sPLS; PLS |
| --- | --- |
| Measure | $R_{\hat{T}T}^2, R_{\hat{T}\hat{U}}^2, \text{TPR}, W^\top \hat{W}$ |
| $p; q$ | 20, 000; 10, 000 |
| relevant $p; q$ | 1000; 500 |
| $N$ | [30, 100, 200, 300, 600] |
| $\sigma_{t_\perp}^2/\sigma_t^2$ | [1/5, 1/3, 1/2, 1, 2, 3, 5] |
| noise level $\alpha$ | [0.1, 0.2, 0.3, 0.4, 0.5, 0.6, 0.7, 0.8, 0.9] |

### Results of simulation study

#### Scenario 1

Fig 1 shows the selection proportion for each relevant group under each noise level. Compared to smaller groups, the proportion for larger groups is higher at low to moderate (*α* < 0.7) noise levels, and shows robustness against increasing noise. When the noise level is very high (*α* > 0.8), the method loses power to detect relevant group of any size, particularly, of larger size. Fig 2 shows the density of the group importance measurement 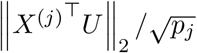 for the first 5 relevant groups with different group sizes under 3 different noise levels. The vertical dotted lines indicate the average threshold given the correct number of relevant groups. Since a group will be selected if exceeds the threshold, the total area on the right side of the threshold under each density curve equals the selection proportion for the corresponding group. The measurement for larger relevant group shows higher precision at all noise levels. The threshold increases along with the noise.

**Figure 1:**
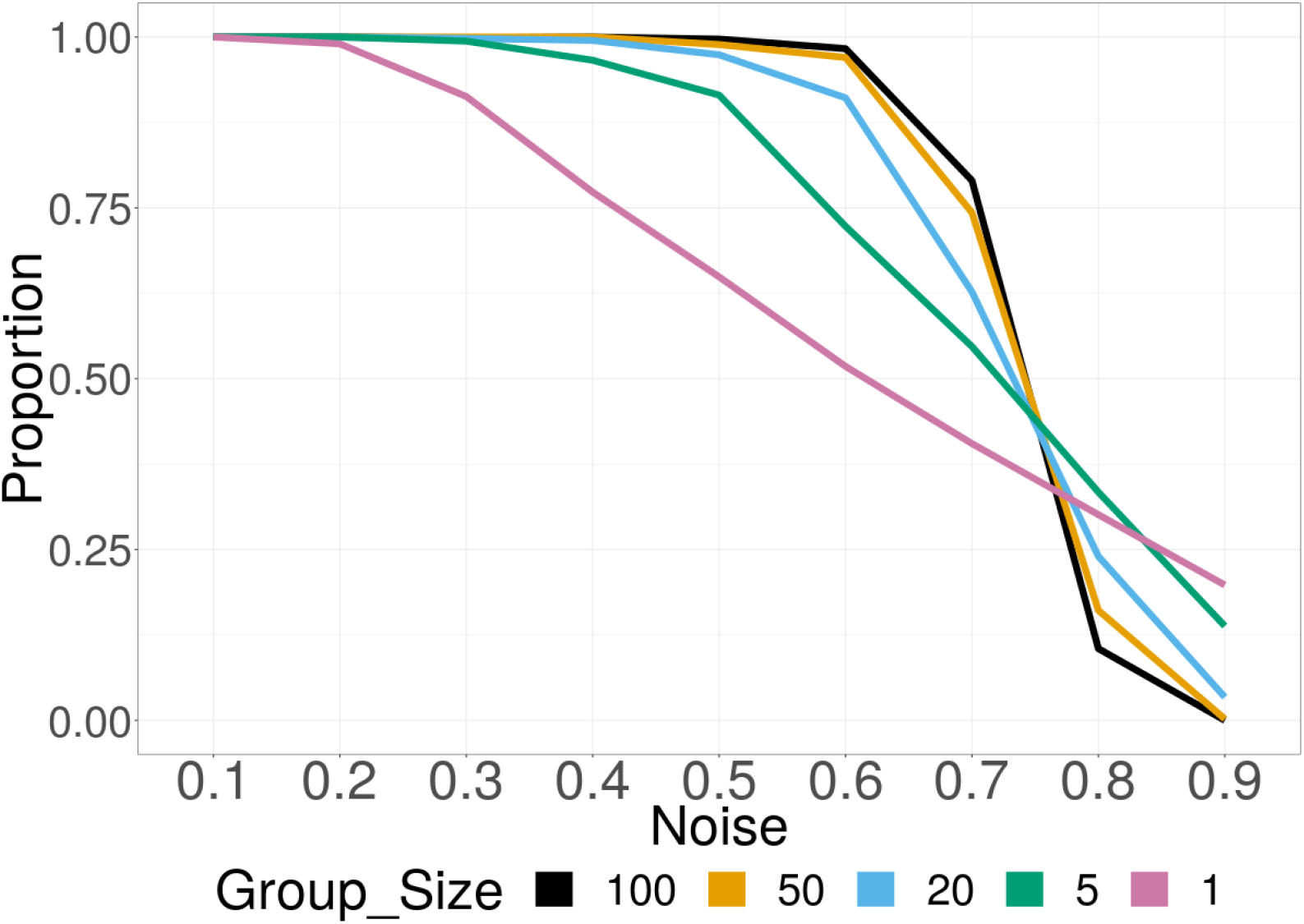
Simulation Scenario 1: Selection proportion of relevant groups with different sizes under varying noise. The proportion for larger groups is higher at low to moderate (*α* < 0.7) noise levels, and shows robustness against increasing noise.

**Figure 2:**
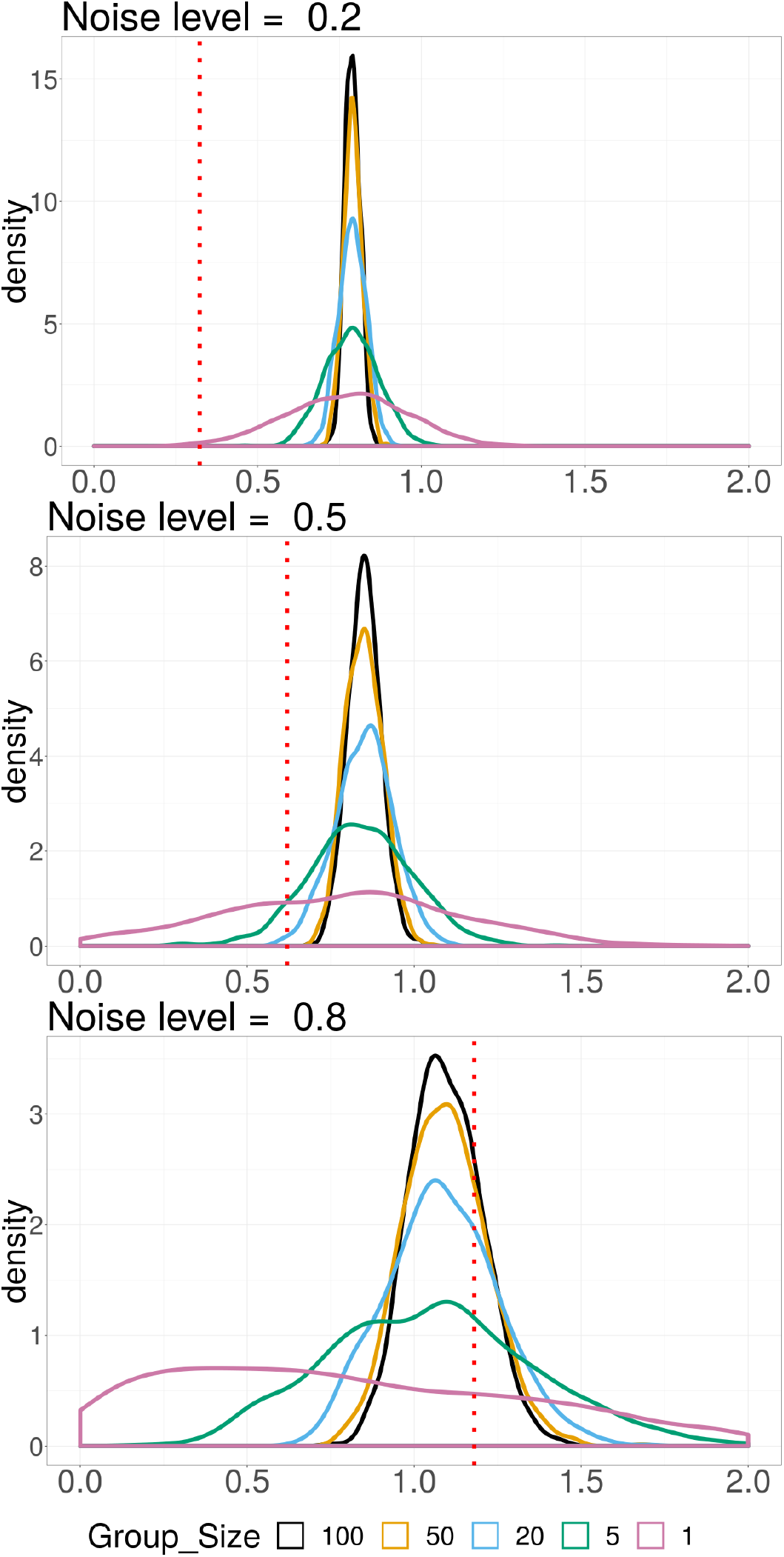
Simulation Scenario 1: Density plot of estimated group importance measurement 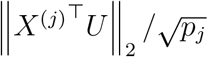 for each group size under 3 different noise levels. The vertical dotted red line is the average threshold. When the measurement of a group is larger than the threshold, the group is selected. The total area on the right side of the threshold under each density curve equals to the selection proportion for the corresponding group. The less the density curve spreads out, the more stable is the estimate.

#### Scenario 2

The performance of the joint score estimation is compared focusing on the difference between methods with orthogonal parts (GO2PLS, SO2PLS, O2PLS) and their counterparts without the “O2” filtering (gPLS, sPLS, PLS). The top row of Fig 3 shows the performance measured by 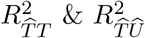under *N* = 30, *α* = 0.1 and varying relative orthogonal signal strength from one fifth to five times of the joint signal. In the left panel, 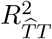 of the various methods is depicted, representing how well the joint component 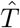 captured the true underlying *T*. Overall, penalized methods performed better than non-penalized ones, especially when the orthogonal variation is relatively small. PLS performed poorly compared to O2PLS, when the orthogonal variation exceeds the joint variation. As the orthogonal variation further increases, performances of sPLS and gPLS deteriorated, while SO2PLS and GO2PLS were less affected. In the right panel, 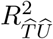 is presented, an estimate of the true parameters 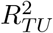, capturing correlation of *T* and *U*. Across different settings, O2PLS-based methods performed better, especially when the orthogonal variation is large.

**Figure 3:**
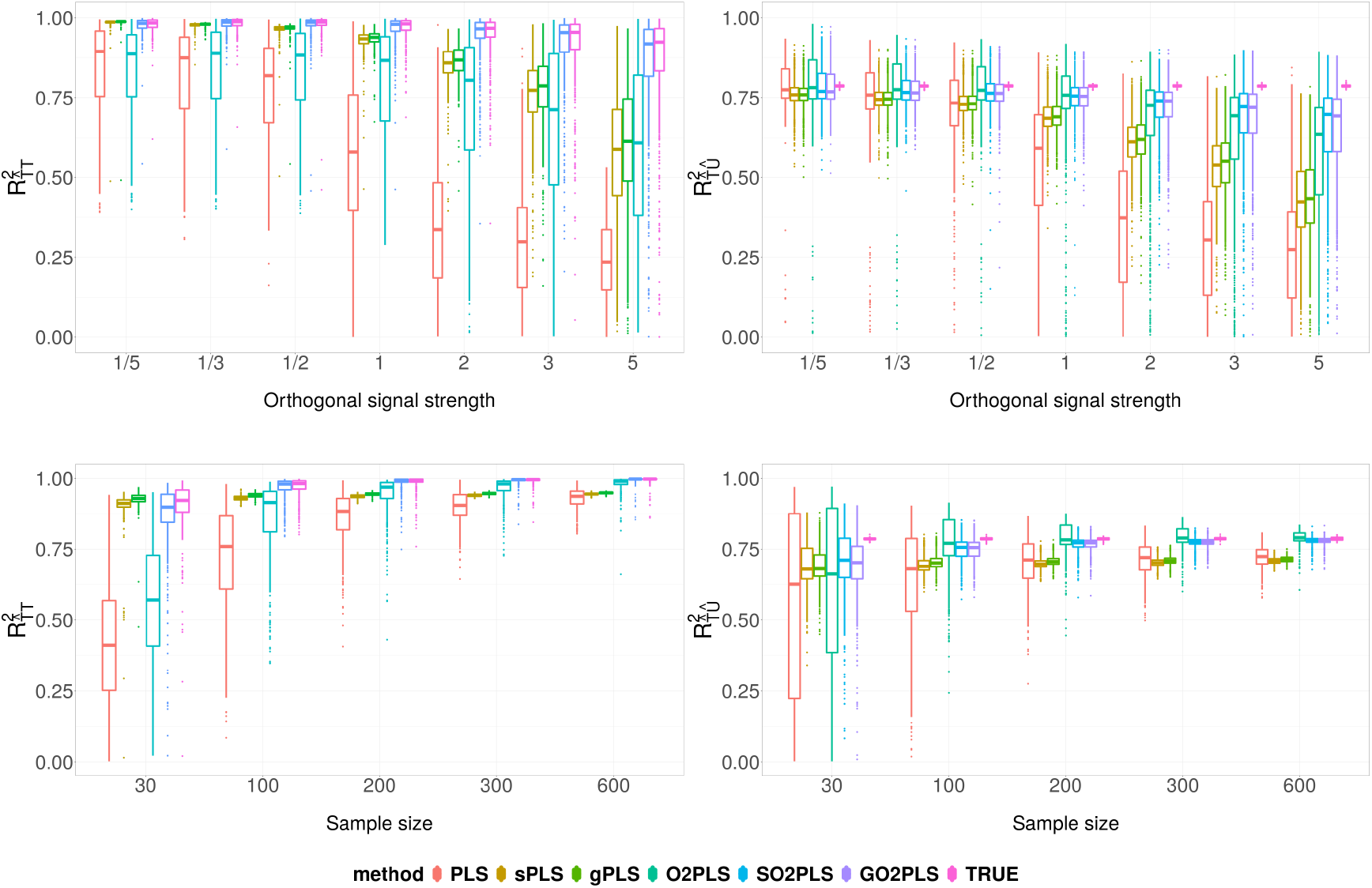
Simulation Scenario 2: comparison of joint score estimation performance, under varying relative orthogonal signal strength (top row), and varying sample size (bottom row). On the Y-axis, 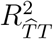 (left) and 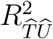 (right) are the coefficient of determination of regressing *T* on 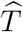, and 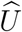 on 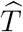, respectively, quantifying the joint score estimation performances. Boxes show the results of 500 repetition.

The bottom row of Fig 3 shows the score estimation performance under fixed relative orthogonal signal strength of 1, *α* = 0.1, and varying sample size *N* from 30 to 600. Penalized methods performed better compared to non-penalized methods in general, when the sample size is small. Regardless of the sample size, O2PLS-based methods outperformed PLS-based methods.

Lastly, we present the results of GO2PLS, SO2PLS, and O2PLS with regard to feature selection and estimation of joint loadings. Results of PLS-based methods are not included since the performances of gPLS, sPLS, and PLS in this regard are very similar to GO2PLS, SO2PLS, and O2PLS, respectively. In Fig 4, the top row shows the TPR and 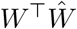 under *N* = 30 and varying noise levels *α* from low to high. At all noise levels, GO2PLS had higher TPR than SO2PLS and O2PLS, and performed robustly against increasing noise. Regarding 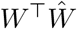, GO2PLS outperformed the other two as well. In the bottom row, when increasing sample size at a fixed noise level of 0.5, the variance appeared to decrease and the performances of all the methods converged. Overall, GO2PLS outperformed others.

**Figure 4:**
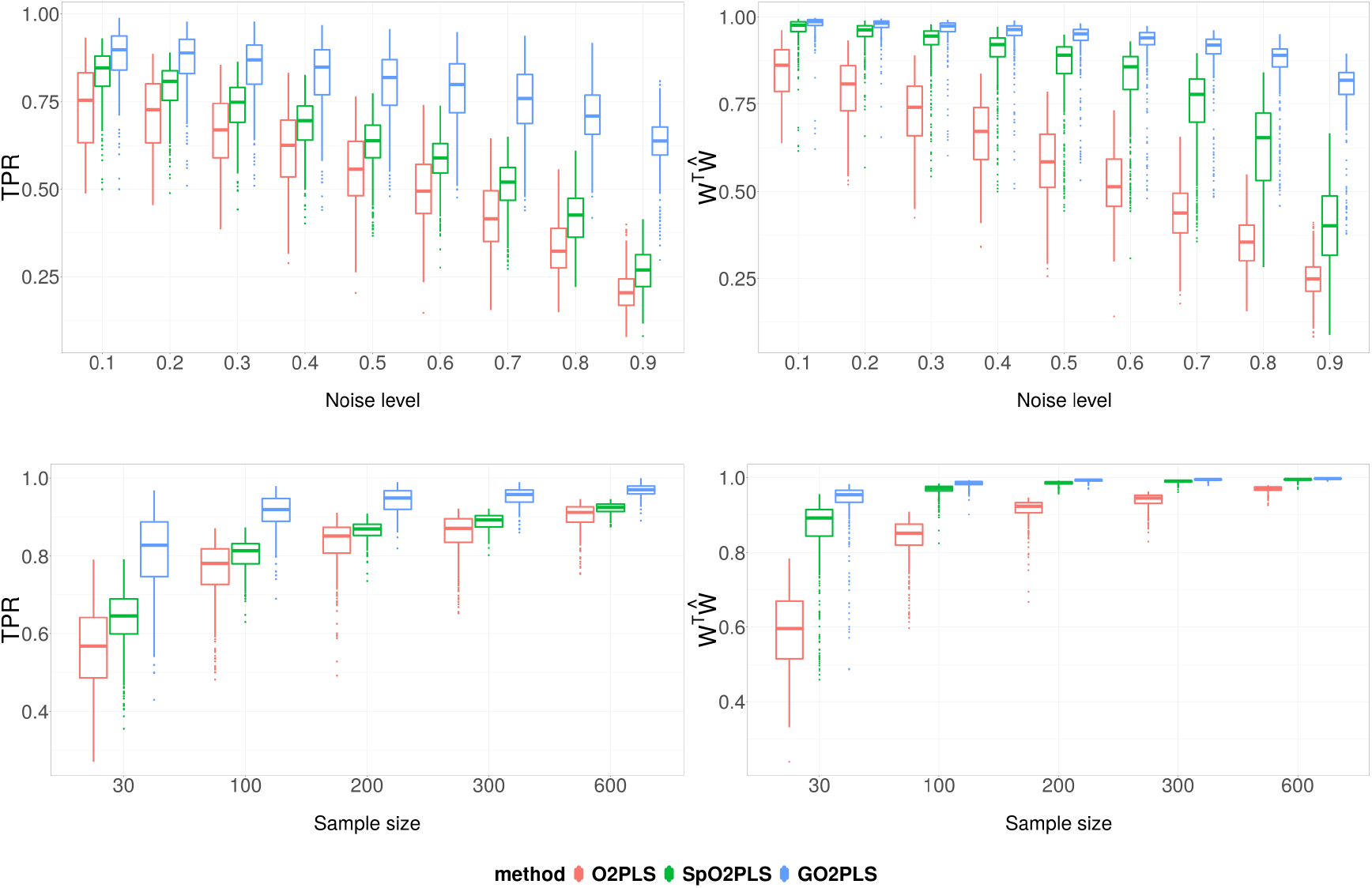
Simulation Scenario 2: comparison of feature selection and joint loading estimation performance, under varying noise level (top row), and varying sample size (bottom row). On the Y-axis are the True Positive Rate (left) and 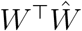 (right), which is the cosine of the angle between the estimated loading vector 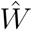 and the true one *W*. Boxes show the results of 500 repetition.

### Application to data

We demonstrate SO2PLS and GO2PLS on datasets from two distinct studies. In the TwinsUK study, our aim is to integrate methylation and glycomics data and identify important groups of CpG sites underlying glycosylation. In the CVON-DOSIS study, we integrate regulomics and transcriptomics data and select a subset of genes and regions that drive their relationship.

### TwinsUK study

We performed GO2PLS on the data with 1 joint, no methylation-orthogonal, and 3 glycomics-orthogonal components based on 5-fold cross-validation. We set the sparsity parameters to select the top 100 groups in the methylation and kept all the 22 glycan variables. The selected CpG groups from GO2PLS were mapped to their targeted genes for interpretation.

We performed gene set enrichment analyses on the selected genes using the ToppGene Suite [6]. The results appeared to be related to immune response. We listed the most significant molecular function, biological process, and pathway in Table 3 (the full list of significant results can be found in Additional file 2).

**Table 3:** TwinsUK study: top results of gene set enrichment analysis.

| GO2PLS | Name | pValue | FDR B&H |
| --- | --- | --- | --- |
| GO: Molecular Function | peptide antigen binding | 1.42E-06 | 5.45E-04 |
| GO: Biological Process | homophilic cell adhesion via plasma membrane adhesion molecules | 1.82E-10 | 4.20E-07 |
|  | cell-cell adhesion via plasma-membrane adhesion molecules | 5.46E-10 | 6.28E-07 |
|  | cell-cell adhesion | 3.43E-07 | 2.63E-04 |
|  | interferon-gamma-mediated signaling pathway | 1.01E-05 | 5.83E-03 |
| Pathway (Source: KEGG) | Viral myocarditis | 8.00E-08 | 9.60E-06 |
|  | Staphylococcus aureus infection | 1.32E-06 | 7.92E-05 |
|  | Allograft rejection | 3.77E-06 | 1.51E-04 |
|  | Graft-versus-host disease | 5.54E-06 | 1.66E-04 |
|  | Type I diabetes mellitus | 7.05E-06 | 1.69E-04 |
|  | Autoimmune thyroid disease | 2.00E-05 | 3.66E-04 |
|  | Rheumatoid arthritis | 2.14E-05 | 3.66E-04 |

**Table 4:**
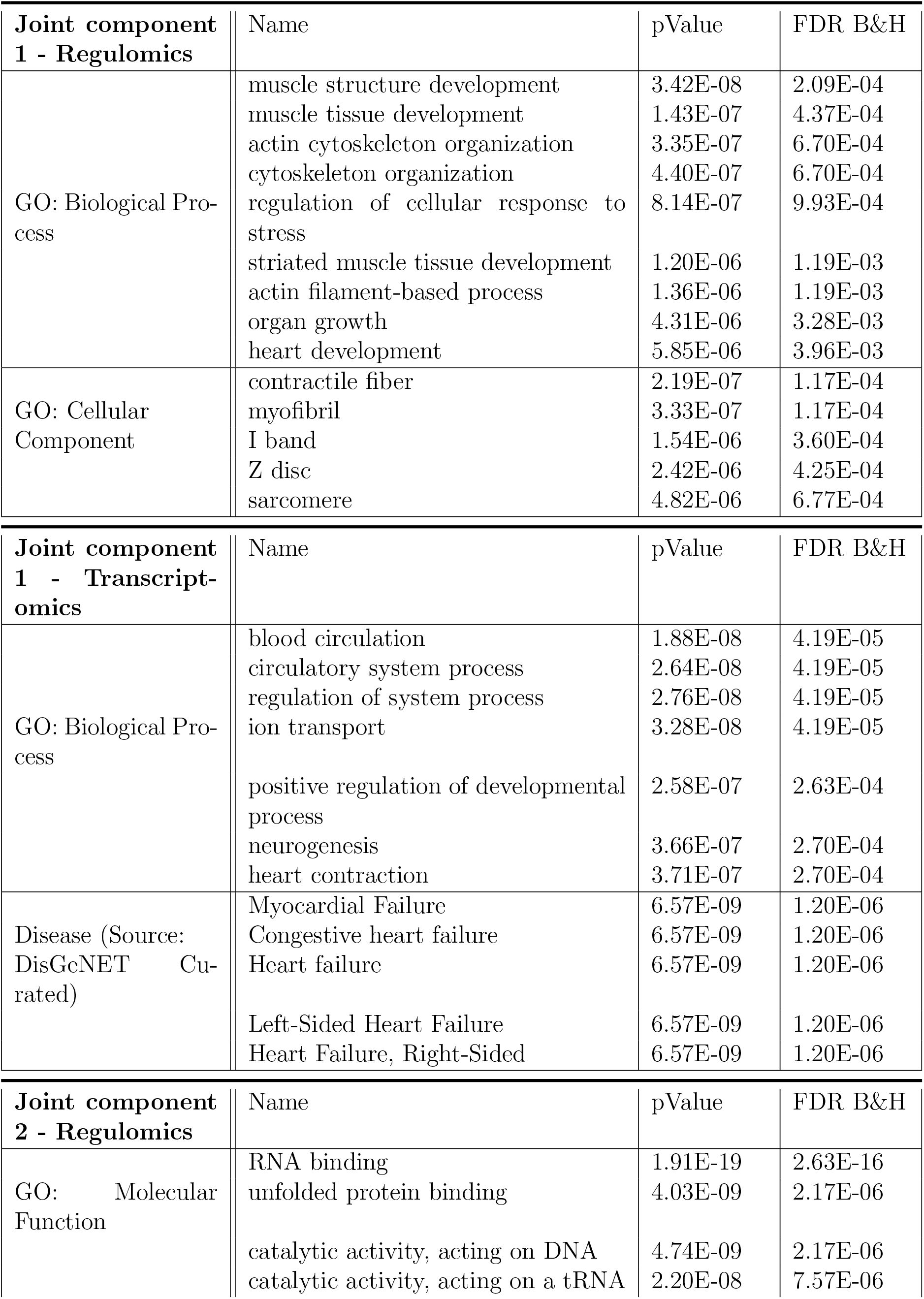
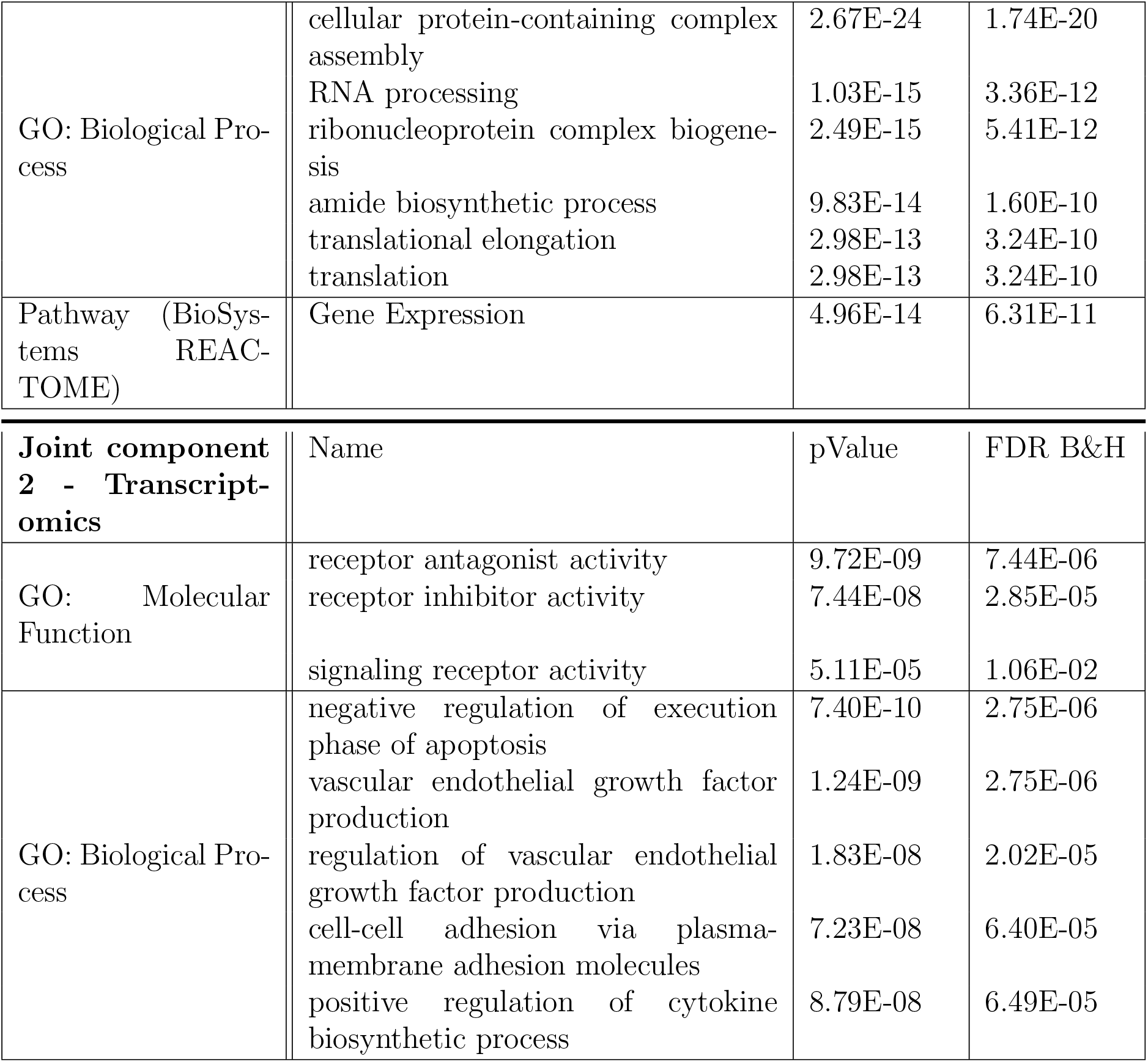
CVON-DOSIS study: Gene set enrichment analysis results.

| **Joint component 1 - Regulomics** | Name | pValue | FDR B&H |
| --- | --- | --- | --- |
| GO: Biological Process | muscle structure development | 3.42E-08 | 2.09E-04 |
|  | muscle tissue development | 1.43E-07 | 4.37E-04 |
|  | actin cytoskeleton organization | 3.35E-07 | 6.70E-04 |
|  | cytoskeleton organization | 4.40E-07 | 6.70E-04 |
|  | regulation of cellular response to stress | 8.14E-07 | 9.93E-04 |
|  | striated muscle tissue development | 1.20E-06 | 1.19E-03 |
|  | actin filament-based process | 1.36E-06 | 1.19E-03 |
|  | organ growth | 4.31E-06 | 3.28E-03 |
|  | heart development | 5.85E-06 | 3.96E-03 |
| GO: Cellular Component | contractile fiber | 2.19E-07 | 1.17E-04 |
|  | myofibril | 3.33E-07 | 1.17E-04 |
|  | I band | 1.54E-06 | 3.60E-04 |
|  | Z disc | 2.42E-06 | 4.25E-04 |
|  | sarcomere | 4.82E-06 | 6.77E-04 |
| **Joint component 1 - Transcriptomics** | Name | pValue | FDR B&H |
| GO: Biological Process | blood circulation | 1.88E-08 | 4.19E-05 |
|  | circulatory system process | 2.64E-08 | 4.19E-05 |
|  | regulation of system process | 2.76E-08 | 4.19E-05 |
|  | ion transport | 3.28E-08 | 4.19E-05 |
|  | positive regulation of developmental process | 2.58E-07 | 2.63E-04 |
|  | neurogenesis | 3.66E-07 | 2.70E-04 |
|  | heart contraction | 3.71E-07 | 2.70E-04 |
| Disease (Source: DisGeNET Curated) | Myocardial Failure | 6.57E-09 | 1.20E-06 |
|  | Congestive heart failure | 6.57E-09 | 1.20E-06 |
|  | Heart failure | 6.57E-09 | 1.20E-06 |
|  | Left-Sided Heart Failure | 6.57E-09 | 1.20E-06 |
|  | Heart Failure, Right-Sided | 6.57E-09 | 1.20E-06 |
| **Joint component 2 - Regulomics** | Name | pValue | FDR B&H |
| GO: Molecular Function | RNA binding | 1.91E-19 | 2.63E-16 |
|  | unfolded protein binding | 4.03E-09 | 2.17E-06 |
|  | catalytic activity, acting on DNA | 4.74E-09 | 2.17E-06 |
|  | catalytic activity, acting on a tRNA | 2.20E-08 | 7.57E-06 |
| GO: Biological Process | cellular protein-containing complex assembly | 2.67E-24 | 1.74E-20 |
|  | RNA processing | 1.03E-15 | 3.36E-12 |
|  | ribonucleoprotein complex biogenesis | 2.49E-15 | 5.41E-12 |
|  | amide biosynthetic process | 9.83E-14 | 1.60E-10 |
|  | translational elongation | 2.98E-13 | 3.24E-10 |
|  | translation | 2.98E-13 | 3.24E-10 |
| Pathway (BioSystems REACTOME) | Gene Expression | 4.96E-14 | 6.31E-11 |

| **Joint component 2 - Transcriptomics** | Name | pValue | FDR B&H |
| --- | --- | --- | --- |
| GO: Molecular Function | receptor antagonist activity | 9.72E-09 | 7.44E-06 |
|  | receptor inhibitor activity | 7.44E-08 | 2.85E-05 |
|  | signaling receptor activity | 5.11E-05 | 1.06E-02 |
| GO: Biological Process | negative regulation of execution phase of apoptosis | 7.40E-10 | 2.75E-06 |
|  | vascular endothelial growth factor production | 1.24E-09 | 2.75E-06 |
|  | regulation of vascular endothelial growth factor production | 1.83E-08 | 2.02E-05 |
|  | cell-cell adhesion via plasma-membrane adhesion molecules | 7.23E-08 | 6.40E-05 |
|  | positive regulation of cytokine biosynthetic process | 8.79E-08 | 6.49E-05 |

### CVON-DOSIS study

We applied SO2PLS on the regulomics and transcriptomics datasets, with 2 joint and 1 orthogonal components for each omics dataset. In each pair of the joint components, 1000 regulomics and 500 transcriptomics variables were selected. We then further identified the genes corresponding to the promoter regions where the selected 1000 histone modification locates (using ± 10K window from the transcription start site of the gene). These genes are of interest since they are likely to be related to epigenetic regulation of gene expression. Genes corresponding to the selected transcripts were also identified. These gene sets identified from each joint component of the two omics data were investigated separately using gene set enrichment analysis. The top results were listed in Table 2. The GO analysis of the selected genes and regions showed terms related to HCM that were also found previously [11]. Due to the presence of the case-control status in both omics levels, we expect the joint components related to the disease. Plotting the joint scores of the two datasets showed a separation between HCM cases and controls (Fig 5). For a comparison of score plots of PCA, PLS, O2PLS, and SO2PLS, please see Additional file 3.

**Figure 5:**
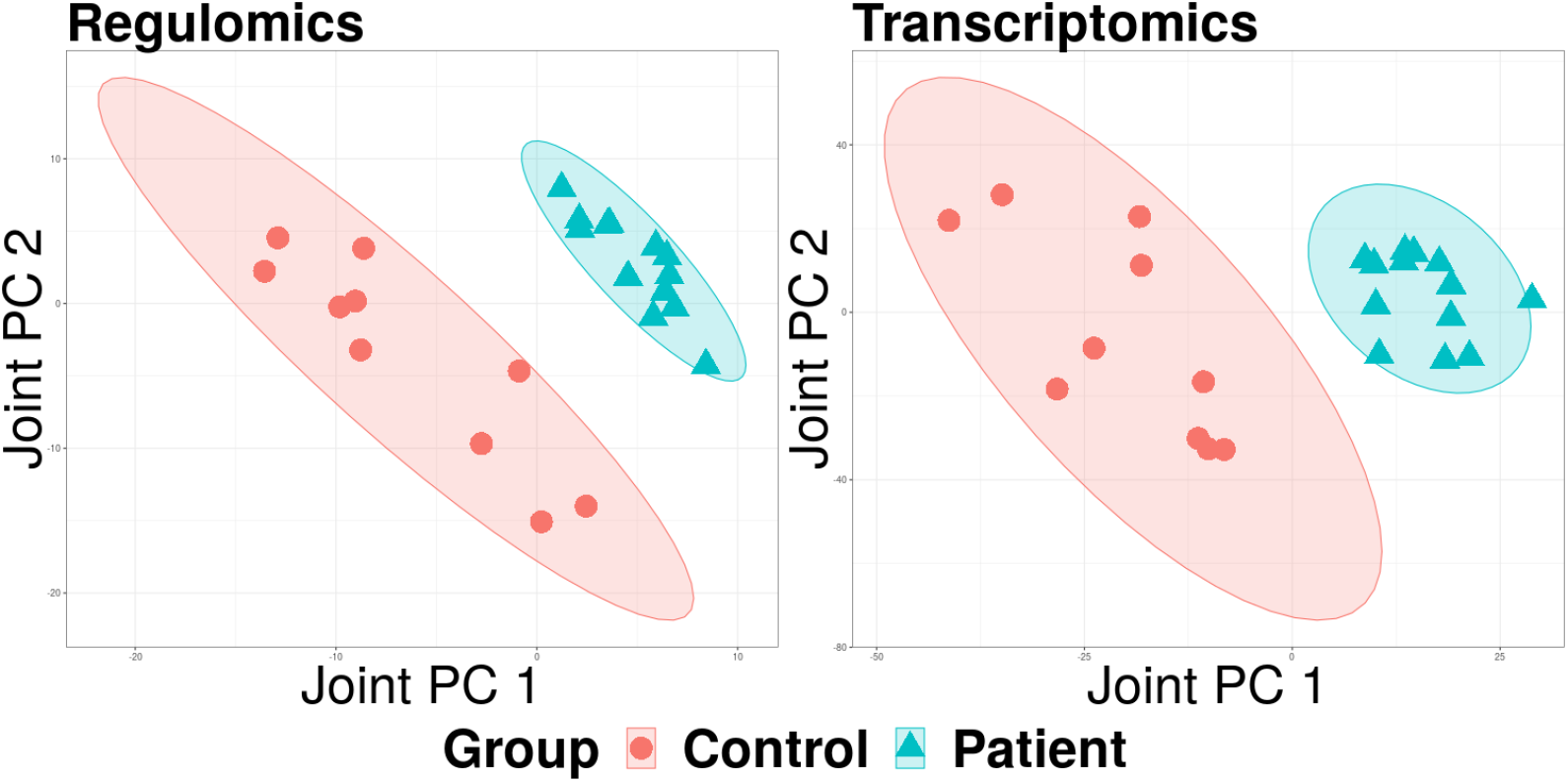
CVON-DOSIS study: SO2PLS joint score plots of regulomics (left) and transcriptomics (right). HCM patients and controls were plotted in different colors. Ellipses are the 95% confidence regions of each group.

## Discussion and conclusion

Statistical integration of two omics datasets is becoming increasingly popular to gain insight into underlying biological systems. O2PLS is a method that integrates two heterogeneous datasets and takes into account omic-specific variation. The resulting joint and specific components are linear combinations of all variables, making interpretation difficult. To introduce sparsity and identify relevant groups, GO2PLS incorporates biological information on group structures to perform group selection in the joint subspace.

Depending on the group size, such an approach may also lead to a higher selection probability of relevant features. We performed an extensive simulation study and showed that O2PLS-based methods generally outperformed PLS-based methods regarding joint score estimation when orthogonal variation was present in the data. Since PLS does not take into account orthogonal parts, the joint components also include part of the orthogonal variation. Further, when the sample size was small or the noise level was high, penalized methods appeared to be much less prone to overfitting than non-penalized methods. This suggests that results based on GO2PLS are likely to be generalizable when applied to new datasets. Concerning feature selection, adding external group information led to higher TPR, and larger groups of relevant features had a higher proportion of being detected under a moderate noise level. We then applied GO2PLS to the TwinsUK study, where we selected 100 target genes comprising of CpG sites that are most related to IgG glycosylation. The results of the enrichment analysis on the selected genes showed GO-terms involving the immune system in which the IgG glycans play important roles. In the CVON-DOSIS study, we integrated regulomics and transcriptomics and identified 1000 regulatory regions and 500 transcripts, and mapped them to genes. Further analysis of the selected gene sets showed enrichment for terms related to heart muscle diseases. Moreover, the implementation of GO2PLS is computationally fast and memory efficient. It relies on an algorithm based on NIPALS that does not store large matrices of size *p* × *q* when performing the group-penalized optimization. A regular laptop (8G RAM, quad-core 2.6 GHz) was able to run GO2PLS on omics data from both case studies.

The group information should be chosen together with domain experts based on the research question and biological knowledge. For example, in our TwinsUK data application, we aimed to identify the genes comprising of CpG sites, rather than the individual CpG sites. Therefore, we grouped CpG sites in the same genetic region. Furthermore, the biological knowledge that close-by CpG sites tend to function together supported the choice of grouping. Different grouping information leads to a changed definition of groups, consequently the selected groups will have a different interpretation. An extra analysis in the TwinsUK study was performed using another grouping strategy. We grouped 55531 CpG sites that map to the promoter region (0-1500 bases upstream of the transcriptional start site (TSS)) of a gene to 14491 groups based on their targeted genes. We applied GO2PLS and selected 100 groups. Note that the size of these groups was smaller, and many CpG sites in gene bodies are excluded. Enrichment analysis did not result in significant results, supposedly due to weaker aggregated group effects. When the research goal is to identify individual features (e.g., in our CVON-DOSIS data application), or group information is not available, SO2PLS can be used.

In the CVON-DOSIS study, Plotting the first two joint components showed two distinct classes corresponding to the case-control status. This might be expected since the analysis was conditional on case-control status, yielding a correlation between the two omics datasets. This phenomenon is well known in regression analysis of secondary phenotypes [23], but not well studied in PLS type of methods. This is a topic of future research. Often omics data are collected to study their relationship with an outcome variable or to predict an outcome variable. To this end, our approach has to be extended to incorporate the outcome variable. Such an approach might also lead to a more sparse solution since the selected features have to be correlated among the three datasets. Further extensions of GO2PLS are to incorporate more than two omics datasets to represent the actual biological system even better.

Finally, it is possible to extend the GO2PLS algorithm to a probabilistic model. Extending latent variable methods to probabilistic models is not new. PCA was extended to Probabilistic PCA in [4], and PPLS [10] was proposed to provide a probabilistic framework for PLS. It has been shown that the probabilistic counterpart has a lower bias in estimation and is robust to non-normally distributed variables [10]. More importantly, the probabilistic model will allow statistical inference, making it possible to interpret the relevance and importance of features in the population, and facilitating follow-up studies. These extensions of GO2PLS will be suited for various studies with more complicated designs.

To conclude, GO2PLS estimates joint latent components that represent underlying systems by integrating two omics data while taking into account the heterogeneity between different omics levels. It incorporates external information on group structures to perform group selection, leading to better interpretation.

## Declarations

### Ethics approval and consent to participate

Not applicable

### Consent for publication

Not applicable

### Availability of data and material

The R scripts and functions for GO2PLS are publicly available in the OmicsPLS R package https://cran.r-project.org/package=OmicsPLS and can be installed in R via install.packages(“`OmicsPLS`”).

Because of the sensitive nature of the data collected for the CVON-DOSIS study, requests to access the dataset from qualified researchers trained in human subject confidentiality protocols may be sent to the corresponding authors.

Individual level methylation and glycomics data from TwinsUK are not permitted to be shared or deposited due to the original consent given at the time of data collection. However, access to methylation and glycomics data can be applied for through the TwinsUK data access committee. For information on access and how to apply, visit http://www.twinsuk.ac.uk/data-access/submission-procedure/.

## Authors’ contributions

ZG performed the mathematical work, simulation study, data anaylysis, and wrote the manuscript. JH and H-WU provided the underlying idea of the methods. SB supported the programming. All authors contributed to the discussion of the methods, simulation and data analysis. All authors read and approved the final manuscript.

## Acknowledgements

The authors would like to thank M. Harakalova, and M. Mokry from the Dept. of Cardiology, UMC Utrecht for providing the CVON-DOSIS data and discussion on the analysis of the CVON-DOSIS datasets. We thank M. Michels and J. van der Velden for providing the HCM tissues, the biobank of UMC Utrecht, the biobank of the Washington University School of Medicine, and the Sydney Heart Bank for providing non-failing donor tissue.

This work has received support from the EU/EFPIA Innovative Medicines Initiative 2 Joint Undertaking BigData@Heart grant (116074).

TwinsUK is funded by the Wellcome Trust, Medical Research Council, European Union, Chronic Disease Research Foundation (CDRF), Zoe Global Ltd and the National Institute for Health Research (NIHR)-funded BioResource, Clinical Research Facility and Biomedical Research Centre based at Guy’s and St Thomas’ NHS Foundation Trust in partnership with King’s College London.

## Additional Files

### Additional file 1

The details of solving the optimization problem (8).

### Additional file 2

The full lists of significant results of gene set enrichment analyses in the TwinsUK study and the CVON-DOSIS study.

### Additional file 3

Additional analysis of the CVON-DOSIS datasets, where score plots of PCA, PLS, O2PLS, SO2PLS are shown and compared.

