## Additional file 3 for "Statistical Integration of Two Omics Datasets Using GO2PLS"

### Supplementary Material for: Statistical Integration of Two Omics Datasets Using Group Sparse O2PLS

#### Additional analysis of CVON-DOSIS study

Due to the case-control study design, we expected that the first few principal components of both datasets that explain most of the variance in the data are related to the disease status. We performed PCA on each dataset separately and plotted the scores of each data (Fig 1a) with different colors for the patients and controls. Though the 95% confidence regions of the two groups overlap, the scores of both data separated the groups quite well. We further investigated if the joint components that explain the covariance between the datasets were also related to the disease. We integrated the two omics data using PLS (2 joint components, Fig 1b), O2PLS (2 joint and 1 orthogonal components for each omics dataset, Fig 1c), and SO2PLS (Fig 1d). The group separation by the first 2 joint components appeared to be clearer comparing to PCA, especially when the data-specific variation was taken into account (i.e., in O2PLS and SO2PLS). More research is needed to quantify the performance of group separation and compare across methods.

The methods we applied are all unsupervised. It is interesting to incorporate the disease status in the model. It is our future work to develop supervised integrative approaches. For more discussions on future directions, please refer to the Discussion section in the article.

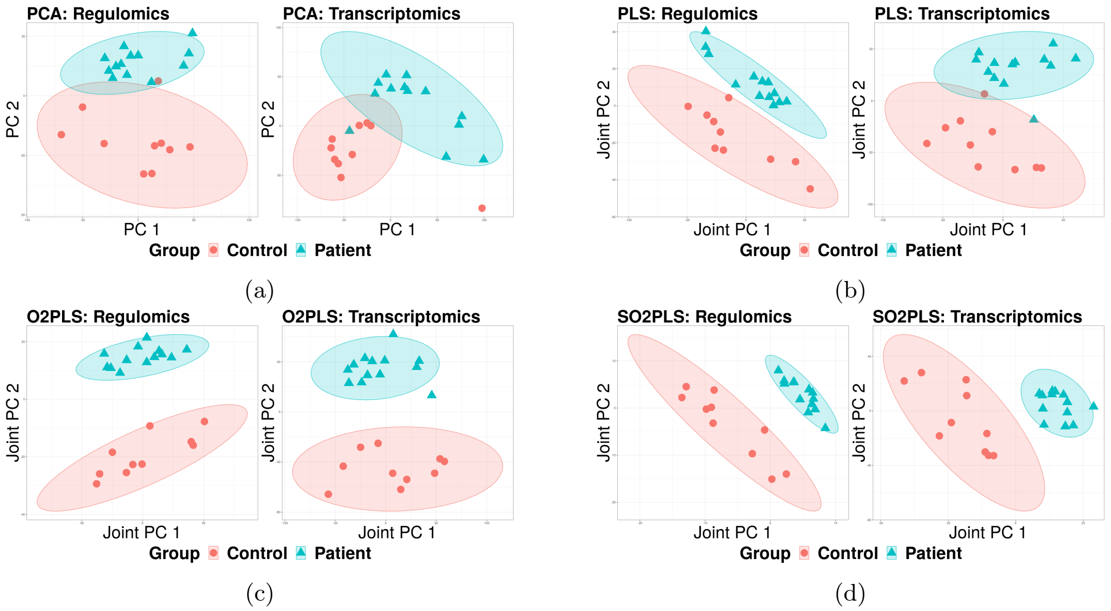


Figure 1: **Score plots of PCA (a), PLS (b), O2PLS (c), and SO2PLS (d); In each subplot, regulomics is on the left, and transcriptomics on the right.** PCA is non-integrative, hence performed on regulomics and transcriptomics separately; PLS, O2PLS, and SO2PLS were applied on the two omics datasets jointly. HCM patients and controls were plotted in different colors. Ellipses are the 95% confidence regions of each group.
