## Additional file 1 for "Statistical Integration of Two Omics Datasets Using GO2PLS"

### Supplementary Material for: Statistical Integration of Two Omics Datasets Using Group Sparse O2PLS

#### Solving the Optimization problem of GO2PLS

The optimization problem of GO2PLS is block multi-convex. It can be solved by optimizing one block at a time, holding the others fixed [1]. The optimization problem for $w_k^{(j)}$ is:

$$\min_{w_k^{(j)}} \left\{ -c_k^\top \tilde{Y}_k^\top \tilde{X}_k^{(j)} w_k^{(j)} + \lambda_w \sqrt{p_j} \left\| w_k^{(j)} \right\|_2 + \phi_w \left( \left\| w_k^{(j)} \right\|_2^2 - 1 \right) \right\}. \tag{1}$$

We differentiate with regard to $w_k^{(j)}$ and set the derivative to 0, and solve for $w_k^{(j)}$:

$$-\tilde{X}_k^{(j)\top} u + \sqrt{p_j} \lambda_w s_j + 2\phi_w w_k^{(j)} = 0, \tag{2}$$

where $s_j$ is the subdifferential of $\left\| w_k^{(j)} \right\|_2$ that has the following form:

$$s_j = \begin{cases} \frac{w_k^{(j)}}{\left\| w_k^{(j)} \right\|_2}, & \text{if } w_k^{(j)} \neq \mathbf{0} \\ \text{Any vector with } \|\hat{s_j}\|_2 \in [-1, 1], & \text{Otherwise.} \end{cases} \tag{3}$$

Rearranging equation (2), we have

$$w_k^{(j)} = \frac{\tilde{X}_k^{(j)\top} u - \sqrt{p_j} \lambda_w s_j}{2\phi_w} = \begin{cases} \mathbf{0}, & \left\| \tilde{X}_k^{(j)\top} u \right\|_2 \in [-\sqrt{p_j}\lambda_w, \sqrt{p_j}\lambda_w] \\ (\tilde{X}_k^{(j)\top} u - \sqrt{p_j}\lambda_w \frac{w_k^{(j)}}{\left\| w_k^{(j)} \right\|_2})/2\phi_w, & \text{Otherwise.} \end{cases} \tag{4}$$

If $w_k^{(j)} \neq \mathbf{0}$, we have

$$w_k^{(j)} = \frac{\left\| w_k^{(j)} \right\|_2}{2\phi_w \left\| w_k^{(j)} \right\|_2 + \sqrt{p_j}\lambda_w} \tilde{X}_k^{(j)\top} u. \tag{5}$$

By taking $L_2$ norm at both sides of (5), we can solve for $\left\| w_k^{(j)} \right\|_2$. Substituting it back to equation (5), we get

$$w_k^{(j)} = \frac{\left\| \tilde{X}_k^{(j)\top} u \right\|_2 - \sqrt{p_j}\lambda_w}{2\phi_w \left\| \tilde{X}_k^{(j)\top} u \right\|_2} \tilde{X}_k^{(j)\top} u. \tag{6}$$

Combining with the case when $w_k^{(j)} = \mathbf{0}$, we have the general form

$$w_k^{(j)} = \frac{\left(\left\|\tilde{X}_k^{(j)\top} u\right\|_2 - \sqrt{p_j}\lambda_w\right)_+}{2\phi_w \left\|\tilde{X}_k^{(j)\top} u\right\|_2} \tilde{X}_k^{(j)\top} u. \tag{7}$$

Similarly, the solution for $c_k^{(m)}$ can be obtained.
